# Steroid hormone imbalance drives macrophage infiltration and *Spp1*/osteopontin^+^ foam cell differentiation in the prostate

**DOI:** 10.1101/2022.12.13.520247

**Authors:** Petra Popovics, Kegan O. Skalitzky, Elise Schroeder, Asha Jain, Samara V. Silver, Francesca Van Fritz, Kristen S. Uchtmann, Chad M. Vezina, William A. Ricke

## Abstract

Benign Prostatic Hyperplasia (BPH) occurs progressively with aging in men and drives deteriorating symptoms collectively known as Lower Urinary Tract Symptoms (LUTS). Age associated changes in circulating steroid hormones, and prostate inflammation have been postulated in the etiology of BPH/LUTS. The link between hormones and inflammation in the development of BPH/LUTS is conflicting because they may occur independently or as sequential steps in disease pathogenesis. This study aimed to decipher the prostatic immune landscape in a mouse model of lower urinary tract dysfunction (LUTD). Steroid hormone imbalance was generated by the surgical implantation of testosterone (T) and estradiol (E2) pellets to male C57BL/6J mice and gene expression analysis was performed on ventral prostates (VP). These experiments identified an increase in the expression of macrophage markers and *Spp1*/osteopontin (OPN). Localization studies of OPN pinpointed that OPN+ macrophages travel to the prostate lumen and transition into lipid accumulating foam cells. We also observed a significantly increased number of tissue macrophages in the VP which was prevented in OPN knockout (OPN-KO) mice. In contrast, mast cells, but not macrophages, were significantly elevated in the dorsal prostate of T+E2 treated mice which was diminished in OPN-KO mice. Steroid hormone implantation progressively increased urinary frequency, which was ameliorated in OPN-KO mice. Our study underscores the role of age associated steroid hormone imbalances as a mechanism of expanding the prostatic macrophage population, their luminal translocation and foam cell differentiation.

## Introduction

Lower Urinary Tract Symptoms (LUTS) in the aging men, such as nocturia, incomplete emptying and weak stream, commonly develop as a result of benign pathological changes in the prostate. Pathology of male LUTS entails non-malignant growth driven by proliferation or resistance to apoptosis, as well as inflammation, smooth muscle dysfunction and fibrosis collectively known as Benign Prostatic Hyperplasia (BPH) (1, 2). Prostatic histological changes associated with LUTS may already occur in a subset of men in their 30s (≤10%) but become highly prevalent (70%) in the sixth decade of life and moderate-to-severe LUTS affect 33% of men in their 60s deteriorating their quality of life (2). As life expectancy increases, BPH/LUTS is becoming an increasing burden on our healthcare system as shown by the 18% increase in BPH patients that was registered between 2004 and 2013 among Medicare beneficiaries (3). Current medical treatments only focus on smooth muscle dysfunction and proliferation while inflammation and fibrosis in the prostate remain clinically untargeted (1, 4). Moreover, the exact etiology and the order of pathological events in the pathogenesis of BPH/LUTS are also poorly understood (2).

Various groups have suggested that an age-related disturbance in prostatic steroid hormone levels may drive BPH pathology. The blockade of the conversion of testosterone to the more potent 5α-dihydrotestosterone by 5α-reductase inhibitors (5-ARIs) has long been used for the reduction of prostate volume. However, patients often develop resistance and progress to surgery despite 5ARI therapy (5). In addition, serum testosterone levels decrease with age, and testosterone on its own often falls short of inducing BPH in animal models (6). In contrast, estradiol levels are sustained in the prostate throughout life leading to an increasing estradiol-to-testosterone ratio during aging (7). In addition, it has been suggested that a synergistic effect of estradiol and testosterone is required to drive BPH pathology (6). An obesity-driven androgenic to estrogenic switch has also been encountered in the prostate (8). Steroid hormone imbalance also causes urinary voiding dysfunction in a mouse BPH/LUTS model which uses slow-release hormone pellets providing supra-physiological levels of estradiol and close to physiological levels of testosterone suggesting that their synergistic effect is required for the full disease course to develop (9, 10).

There are still conflicting findings on how the inflammatory environment is remodeled during steroid hormone imbalance and how that contributes to prostate fibrosis. Rat prostates were shown with either T-lymphocytic and neutrophilic infiltration (11) or increased number of macrophages (12). Contrasting this, others did not find a change in immune cells, including macrophages, in steroid hormone treated mice (13).

The goal of our study was to assess early molecular and cellular changes in the immune environment that is driven by the changing estradiol-to-testosterone ratio. We identified that steroid hormone imbalance drives a spike in the expression of pro-inflammatory genes *Spp1* (osteopontin) and *Saa1* and macrophages in the ventral prostate and mast cells in the dorsal prostate. We also identified that treatment drives the migration of macrophages to the prostate lumen and their differentiation to lipid-laden foam cells that express high level of osteopontin. Finally, we showed that the loss of osteopontin leads to decreased macrophage and mast cell numbers and reduced collagen expression and proliferation.

## Results

### Steroid hormone imbalance drives gene expression signature of tissue remodeling, macrophage infiltration and non-classical pro-inflammatory genes in the mouse prostate

Our primary goal was to identify key inflammatory elements associated with the early stages of steroid hormone imbalance in the prostate. For this reason, we utilized a well-described model generated by the subcutaneous implantation of slow-release testosterone and estradiol (T+E2) hormone pellets (9) and harvested prostatic tissues two weeks later. We performed a Nanostring analysis by isolating RNA from the mouse ventral prostate lobe which we predicted would harbor the most CD45+ inflammatory cells after exogenous hormone treatment, based on our preliminary results (data not shown). We utilized the mouse fibrosis panel (v2) which can detect the expression of 770 genes related to inflammation, tissue remodeling, damage and proliferation. A heatmap generated by NSolver shows that ventral prostate gene expression patterns segregate by treatment group (Fig. S1). Global significance analysis identified the top pathways that were impacted by steroid hormone imbalance as extracellular matrix (ECM) degradation and synthesis, oxidative stress, PDGF signaling and Th1 differentiation. (Fig. 1A, Table S1).

**Figure 1.**
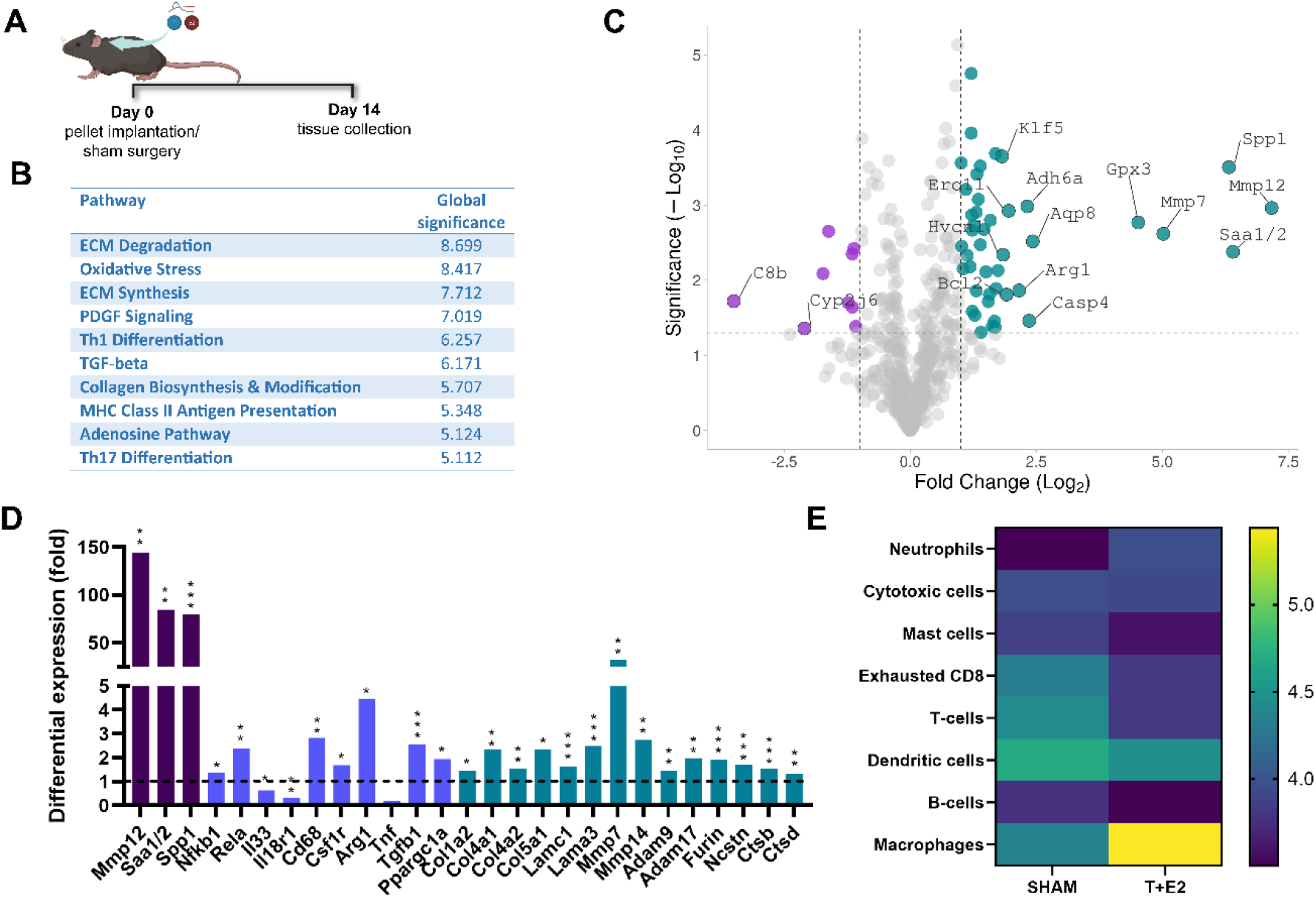
Steroid hormone imbalance promotes gene expression signatures related to inflammation and tissue remodeling in the mouse ventral prostate. Mice were implanted with testosterone and estradiol pellets or received sham surgery and were necropsied 14 days later (A). The NanoString RNA expression technology was used with a mouse fibrotic panel to analyze expressional changes in the ventral prostate lobes. Top ten pathways altered by steroid hormone imbalance included extracellular matrix (ECM) remodeling and oxidative stress amongst inflammatory pathways (B). Top 15 genes were identified by Volcano Plot analysis using VolcaNoseR (C). Selected genes are also depicted in (D) grouped as follows: top three genes, Mmp12, Saa1/2 and Spp1 (dark purple), inflammatory genes/immune cell markers (violet) and extracellular matrix-associated genes (teal, D). Immune cell analysis performed on nCounter Advanced Analysis Software shows the dominance of macrophages in steroid hormone imbalance (E). *: p ≤ 0.05, **: p ≤ 0.01, ***: p ≤ 0.001.

Volcano plot analysis identified *Saa1/2* (serum amyloid A 1 and 2, 84.4-fold), *Mmp12* (matrix metalloproteinase 1, 144.0-fold) and *Spp1* (secreted phosphoprotein 1/ osteopontin, 79.9-fold) as top three upregulated genes (Fig. 1C and 1D). Spp1 and Saa1/2 have important pro-inflammatory roles in part by stimulating the migration of leukocytes (14, 15). In addition, expression of the pro-inflammatory Nuclear factor-κB (NFκB) protein subunit genes, *Nfkb1* and *Rela*, were significantly induced (1.4-fold and 2.4-fold, respectively, Fig. 1E). In contrast, expression levels of major interleukins and *Ccl* or *Cxcl* chemokines were unchanged or were below the level of detection, except *Il33* (driver of Th2 differentiation) and *Il18r* (expressed on T-cells, granulocytes and NK cells), which were significantly reduced (Fig. 1D and Table S2).

Immune cell type analysis predicted a significant increase in macrophages (Fig 1E) in prostates from hormone-treated mice. In addition, macrophage markers *Cd68* (2.8-fold, p=0.008) and *Csf1r* (1.7-fold) and the M2 marker *Arg1* (4.5-fold) were significantly elevated by exogenous hormones whereas the M1 macrophage marker *Tnf* expression fell slightly short of significance for being suppressed (0.2-fold, p=0.053, Fig. 1D and Table S2). Other significantly increased M2 differentiation markers included *Tgfb* (1.35-fold) and *Ppargc1a* (0.954-fold, Table S2) further suggesting the predominance of alternatively activated macrophages.

As suggested by the global significance analysis, various genes related to tissue remodeling were altered by steroid hormone pellets. Matrix structural proteins, including collagens (*Col1a2, −4a1, −4a2 and −5a1*) and basal membrane components (*Lamc1* and *Lama3*) as well as genes involved in ECM turnover, such as matrix metalloproteinases (*Mmp7, −12* and −*14*), “a disintegrin and metalloproteinases” (*Adam9* and −*17*) and other proteases/peptidases (*Furin, Ncstn, Ctsb, Ctsd* and *Ctsl*) were upregulated (Fig. 1D and Table S2). Pathway analysis also indicated changes in oxidative stress-related genes (genes with significant change: *Atp7a, Sod1, P4hb, Ero1l, Cat, Gpx3*, *Aqp8*, *Prdx1*, *Ccs* and *Txn2*, Table S2). Upon further assessment, exogenous steroid hormones significantly reduced several genes encoding proteins of the mitochondrial electron transport chain (*Ndufa1, Ndufb8, Ndufs5, Uqcrfs1*) whereas *Sod1* decreased and *Gpx* elevated (Table S2), potentially as a response to oxidative stress.

### Steroid hormone imbalance drives luminal migration of *Spp1*+ macrophages

Next, we used chromogenic (RNAscope) and fluorescent *in situ* hybridization (FISH), to investigate the tissue localization of the top three elevated genes. *Mmp12* expression was low and was restricted to only a few prostate epithelial cells in the T+E2 group but was absent in sham animals (Fig. 2A). *Saa1/2* expression was highly upregulated in groups of prostate epithelial cells specifically in the T+E2 group, but this induction was sporadic (Fig. 2A). In contrast, *Spp1* expression was highly upregulated, specifically in cells in the prostate lumen (Fig. 2A). To obtain higher sensitivity, we also used FISH for labeling *Spp1* and assessed its co-localization with the macrophage marker, *Cd68* (Fig. 2B). This identified low expression of *Spp1* in epithelial cells and tissue macrophages, but high induction in macrophages that migrated into the lumen in the T+E2 model (Fig. 2B). *Spp1* was also occasionally elevated in unidentified elongated cells in the periglandular smooth muscle layer and had a small elevation in some epithelial regions in response to T+E2 treatment (Fig. 2C). To assess protein levels of SPP1/osteopontin (OPN), we performed immunohistochemistry (IHC) and found that OPN is significantly elevated and localized to the epithelial layer and intraluminal cells as well as to scattered cells in the stroma in the ventral prostate (VP, Fig. 2D and 2E). In contrast, OPN-positive intraluminal cells were not present in the dorsal prostate (DP) and OPN was mainly elevated in cells in the stroma (Fig. 2D and 2F). Upon further investigation, we identified that the steroid hormone imbalance-related luminal macrophage infiltration occurs specifically in the ventral lobe and does not occur in other lobes (data on the anterior and lateral lobes are not shown).

**Figure 2.**
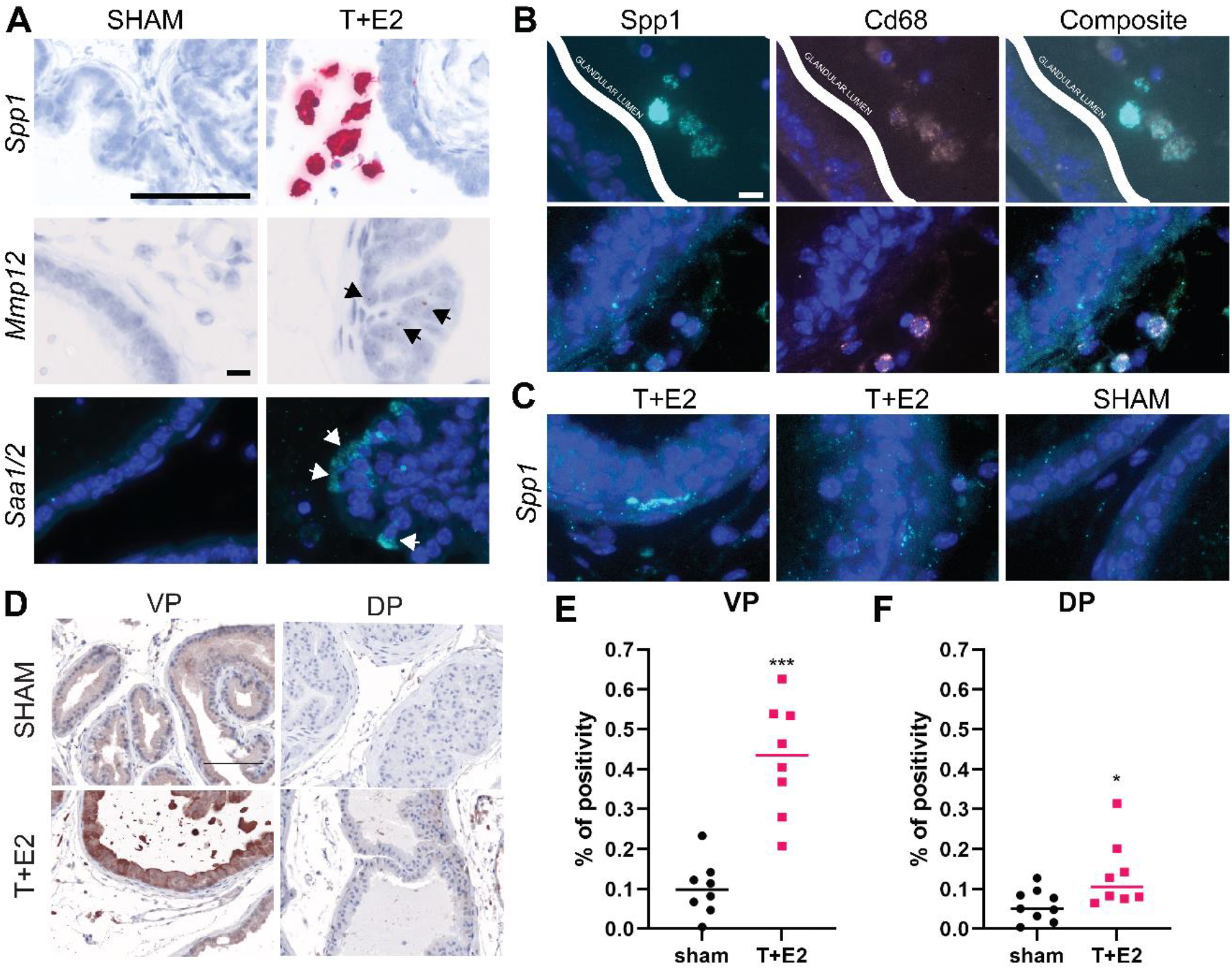
Steroid hormone imbalance drives luminal migration of macrophages and provokes *Spp1* expression in the prostate. Using *in situ* hybridization, *Spp1* was primarily localized to cells in the lumen, *Mmp12* possessed low increase in luminal cells and *Saa1/2* was highly induced in a limited number of epithelial cells (A). *Spp1* showed low expression in the epithelium and in tissue macrophages (co-localization with Cd68), but was highly elevated in macrophages that appeared in the lumen in T+E2 mice (B). Spp1 was also upregulated in a very few cells that localized subepithelially and have slightly higher expression in the epithelium in T+E2 mice compared to sham animals (C). Protein expression of osteopontin (*Spp1* protein product) was highly elevated in the epithelium and some stromal cells in the ventral prostate (VP) but it was only increased in the stroma in the dorsal prostate (DP, D). Quantification of the % of osteopontin positive cells showed significance increase in both the VP and the DP in T+E2 mice (E and F). Significance was calculated by Mann-Whitney test. Scale bars correspond to 100 μm or 10 μm. *: p ≤ 0.05, ***: p ≤ 0.001.

### Luminal macrophages undergo foam cell differentiation

Morphology of luminal macrophages was further investigated using IHC for CD68 (macrophages) or vimentin (labels fibroblasts, astrocytes, endothelial cells and macrophages). This staining helped us to identify the granular appearance which led us to propose that macrophages become lipid-accumulating cells in the lumen, also known as foam cells (Fig 3A). We observed similar morphology in human BPH prostate tissues in Cd68+ macrophages (Fig. 3B). To confirm our hypothesis, we used unfixed frozen sections of mouse ventral prostate and confirmed accumulation of lipid droplets in luminal foam cells using Oil Red O staining (Fig. 3C). The same method on human BPH tissues also confirmed accumulation of lipid-laden foam cells in glandular luminal spaces but also some lipid-dense regions in the epithelial layer (Fig. 3D). Low magnification images of Oil Red O staining showed lipid accumulation in larger nodules (Fig. S2). Using IHC for the M1 macrophage marker, Inducible nitric oxide synthase (iNOS) and the M2 macrophage marker Arginase 1 (ARG1), we identified that luminal macrophages predominantly polarize to M2 macrophages, although M1s were also observed (Fig. 3E and 3F).

**Figure 3.**
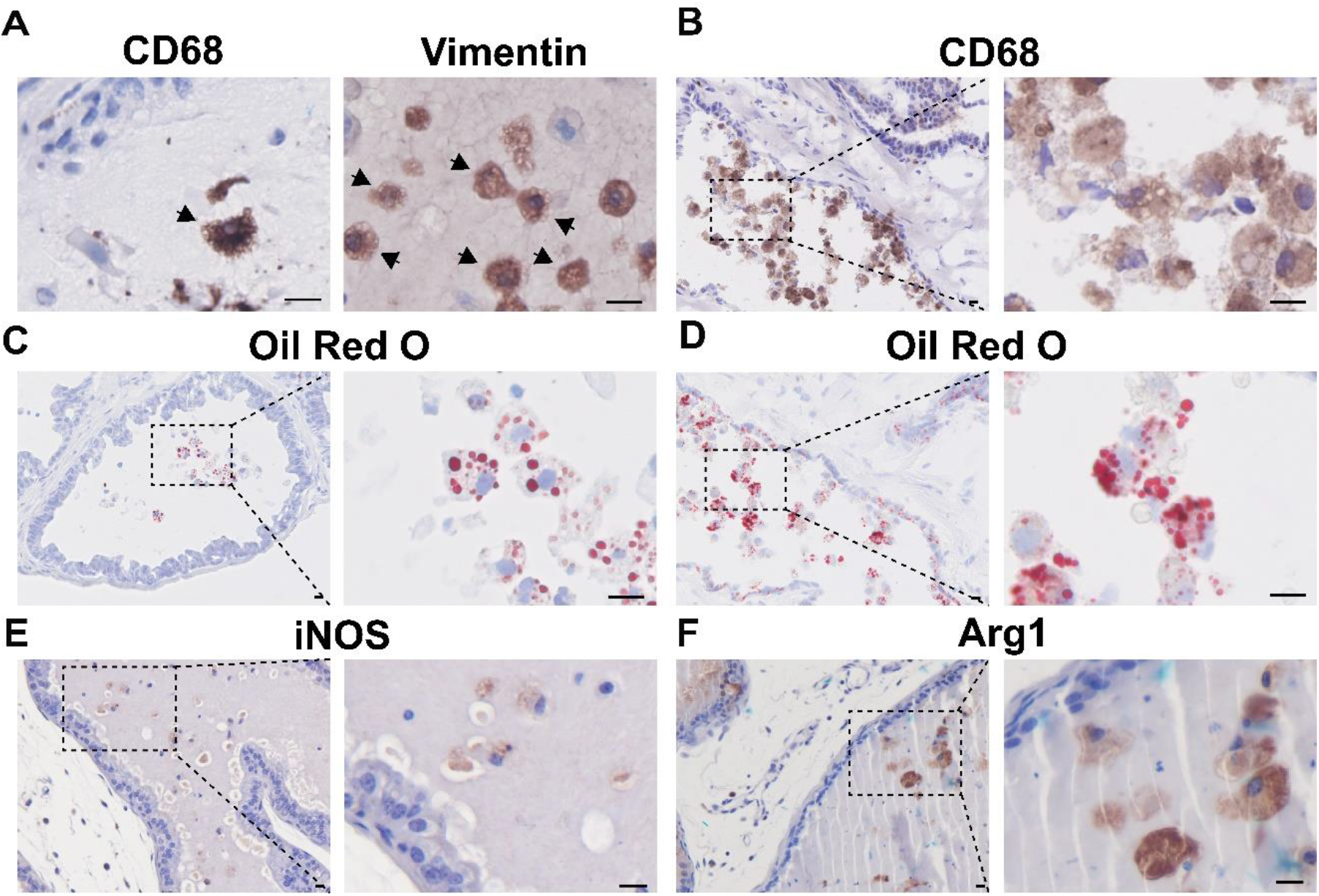
Steroid hormone imbalance instigates foam cell differentiation of luminal macrophages. To study luminal macrophages further, we used immunohistochemical labeling of CD68 (macrophage marker) and Vimentin (expressed in fibroblasts, astrocytes, endothelial cells and macrophages) in T+E2-treated mice. Sham mice contained no luminal macrophages (data not shown). Note the granular appearance of luminal cells with both markers (arrows). CD68+ macrophages can also be found at high numbers in the lumen in human benign prostatic hyperplasia (BPH) tissue (B). Oil Red O staining of mouse T+E2-treated (C) and human BPH specimens (D) identified that the majority of luminal macrophages take up lipids and differentiate into lipid-laden foam cells. Via the detection of iNOS (M1 marker, E) and Arg1 (M2 marker, F), we identified few M1, but more M2 macrophages in the luminal space. Scale bars correspond to 10 μm. Images were taken at 40x magnification.

To better understand the molecular mechanism contributing to increased lipid levels, we assessed genes of lipid synthesis pathways available in our NanoString dataset. We found that the expression of *Fasn* (Fatty Acid Synthase) which encodes a key enzyme of the *de novo* fatty acid synthesis pathway, has significantly increased (1.9-fold, p=0.025) (Table S2).

### Macrophages are the predominant prostatic immune cell type in steroid hormone imbalance

To evaluate the changes in the immune environment in steroid hormone imbalance, as well as to assess the role of OPN in the process, steroid hormone pellet implantation was performed in OPN knockout (OPN-KO) and WT animals for the subsequent analysis of immune cells in the prostate. Two weeks after implantation, CD45+ cells (pan-immune cell marker) increased the most in the ventral lobe of hormone-treated mice (120% increase), but were also elevated in other lobes (Fig. 4, Fig. S3). In contrast, there was no significant change in OPN-KO mice. Six weeks after the procedure, CD45+ cells were significantly increased in the VP in both WT and OPN-KO mice, increased in WT and reduced in OPN-KO AP and were unchanged in the DP and LP. This indicates that an early immune response is most pronounced in the VP, and it is hindered by the systemic loss of OPN.

**Figure 4.**
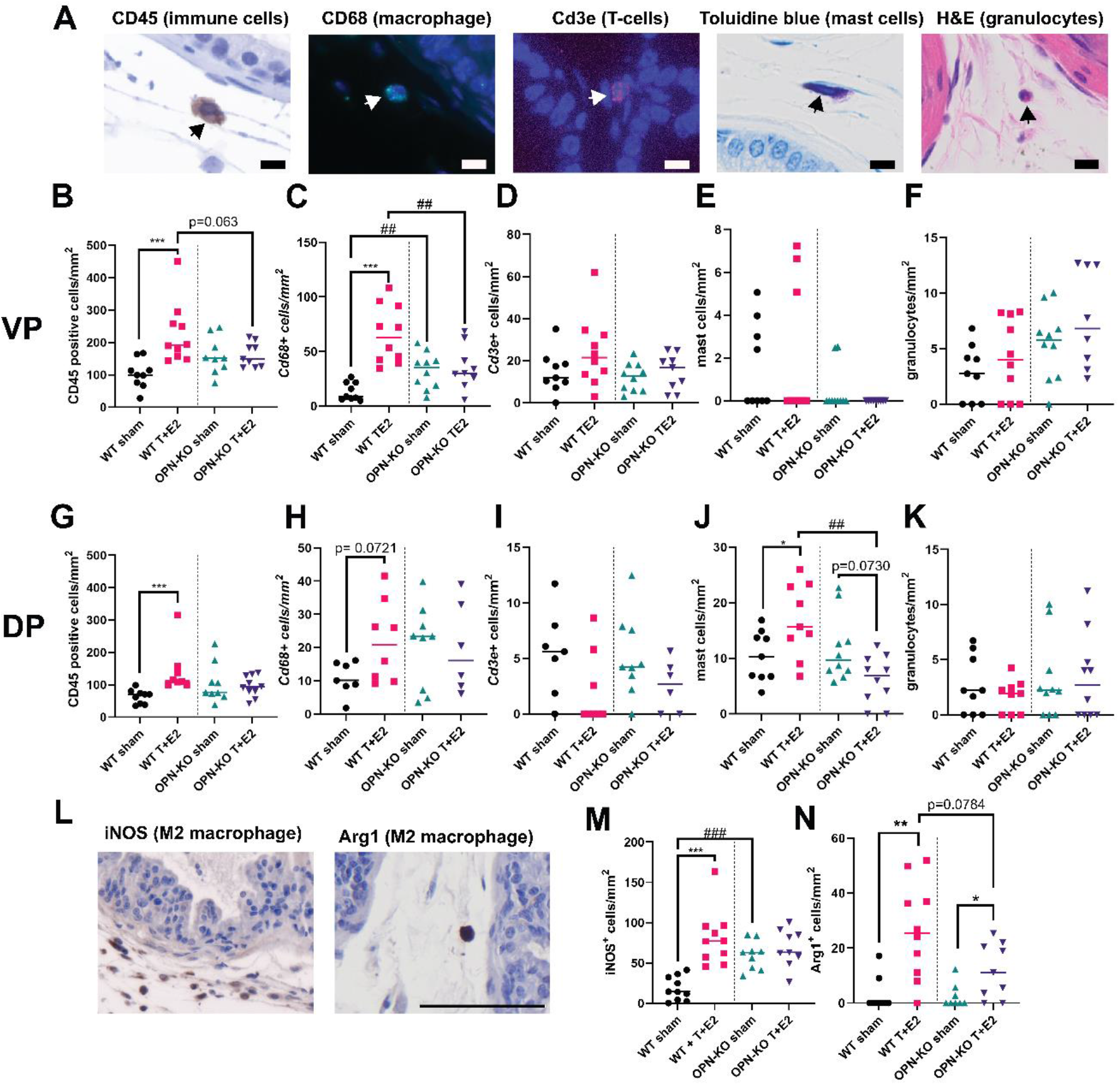
Macrophage and mast cell numbers increase in response to testosterone and estradiol (T+E2) treatment in WT but not in OPN-KO mice. Panel (A) shows example images of staining performed in this study. Immunohistochemical staining of CD45 was performed to identify immune cells, fluorescent in-situ hybridization (FISH) of *Cd68* and *Cd3e* was performed to identify macrophages and T-cells, respectively, toluidine blue staining was used to identify mast cells, and hematoxylin and eosin staining was analyzed to identify granulocytes based on cytoplasmic staining and nuclear morphology. A significant increase in immune cells and macrophages was found in WT but not in OPN-KO mice (B-F). Panels (B), (C), (D), (E) and (F) show CD45, *Cd68, Cd3e*, mast cell, and granulocyte cell counts in the ventral prostate (VP) two weeks after sham or T+E2 pellet surgery. A significant increase in mast cells was found in WT but not in OPN-KO mice (G-K). Panels (G), (H), (I), (J) and (K) show CD45, *Cd68, Cd3e*,mast cell, and granulocyte cell counts in the dorsal prostate (DP) two weeks after sham or T+E2 pellet surgery. FISH for *Cd19* to identify B-cells was performed but was not quantified due to the limited number of positive cells in the prostate in this model. Panel (L) shows representative images of IHC staining for M1 marker iNOS and M2 marker Arg1 in WT T+E2 mice. Quantification of cells show a significant upregulation of iNOS+ cells in WT T+E2, but not in OPN-KO T+E2 mice compared to sham controls (M). Arg1+ cells were upregulated in both WT and OPN-KO mice treated with T+E2. Images (6/tissue) were taken at 40x magnification. Scale bars correspond to 10 μm. All cells were counted manually and were normalized to tissue area as cells/mm^2^. Luminal immune cells were excluded from quantification. Significance was tested by Mann-Whitney test. WT sham vs. WT T+E2 or OPN-KO sham vs. OPN-KO T+E2: *: p≤0.05; ***: p≤0.001. WT T+E2 vs. OPN-KO T+E2 or WT sham vs. OPN-KO sham: ##: p≤0.01; ***: p≤0.001. WT sham vs. OPN-KO T+E2 comparison is not shown.

Next, we narrowed down the scale of investigation to the VP and the DP and utilized FISH, toluidine blue and H&E staining and quantification strategies to identify changes in specific immune cells or groups. We found that steroid hormone imbalance causes a significant increase in CD68+ macrophage numbers in the VP and also a near-significant elevation in the DP (p=0.0721). Interestingly, basal macrophages in the VP were also significantly elevated in OPN-KO sham vs WT sham animals. In contrast, VP macrophage densities were lower in OPN-KO T+E2 vs. WT mice (p≤0.01). These calculations excluded luminal macrophages, which appeared in both WT and OPN-KO T+E2 VPs in similar numbers but were absent in the DP and were not counted. In the DP (but in in the VP), total number of mast cells are also significantly elevated in WT but not in OPN-KO in response to steroid hormone imbalance. In contrast, activation of mast cells is not significantly increased in the DP (Fig. S4), but reduced by hormones in OPN-KO T+E2 mice. We did not find a significant increase in T-cells (*Cd3e*) or granulocytes (H&E, Fig. 4), and were unable to detect B-cells (*Cd19*) in any of the conditions (data not shown).

To identify the predominant tissue macrophage polarization state in the VP on week two, we performed IHC for iNOS (M1 marker) and Arg1 (M2 marker) (Fig. 4 L, M and N). We found a 4.4-fold elevation in iNOS^+^ cells in WT T+E2, but no change in OPN-KO T+E2 mice, compared to their respective sham controls. We also identified a significant elevation in iNOS^+^ cells in OPN-KO sham vs. WT sham ventral prostates. The proportion of ARG1^+^ M2 macrophages was much lower compared to M1 macrophages (26.3 cells/mm^2^ vs 80.9 iNOS^+^ cells/mm^2^ in WT T+E2), but was significantly upregulated in both WT and OPN-KO T+E2 mice vs. sham controls. These results identify macrophages, with a primarily M1 polarization state, as the predominant immune cells responding to steroid hormone imbalance in the VP, whereas the increased mast cell presence may be primarily important in the DP in this model.

### Loss of osteopontin delays the onset of urinary dysfunction and suppresses prostatic fibrosis and proliferation

T+E2 mice develop urinary dysfunction which manifests, in part, as increased urination frequency and decreased voiding volume/void (9). To assess whether the loss of OPN impacts urinary function in T+E2 mice, we performed weekly void spot assays from week one to five after pellet implantation and analyzed the number of voiding events (frequency). WT mice had increased spot numbers starting on week two which continued to increase throughout the experiment. In contrast, OPN-KO T+E2 mice had significantly fewer void spots until week four (Fig. 5A) demonstrating that the loss of OPN delays the onset of steroid hormone imbalance-induced urinary dysfunction.

**Figure 5.**
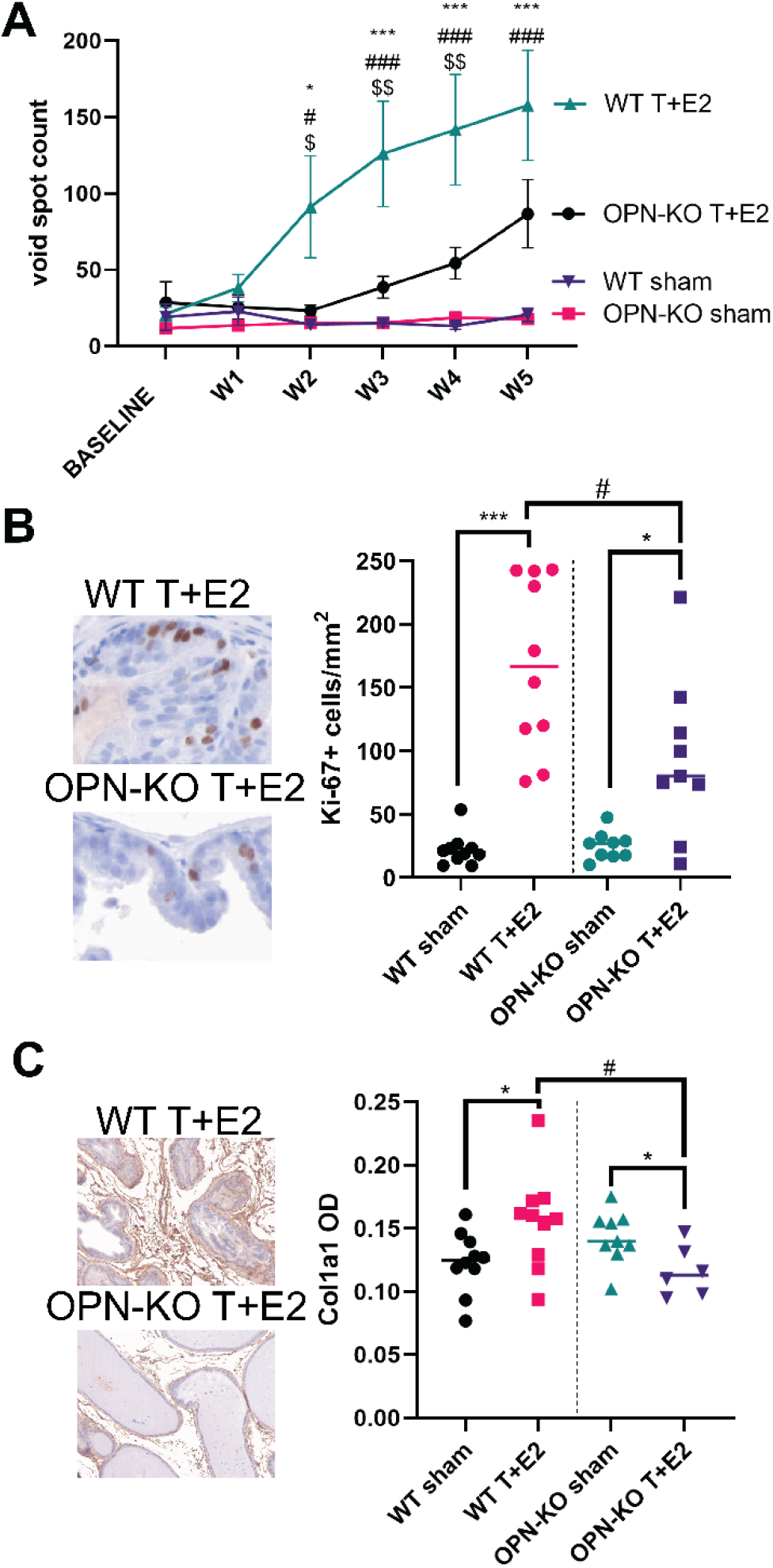
Loss of OPN delays the onset of urinary dysfunction, reduces proliferation and prevents prostatic fibrosis induced by steroid hormone imbalance. Urinary voiding frequency induced by T+E2 pellet implantation was ameliorated by the loss of OPN until week four (A). Ki-67-proliferation marker was significantly increased in both WT and OPN-KO T+E2 ventral prostates compared to sham animals, but were reduced in OPN-KO T+E2 mice compared to WT T+E2 (B). Collagen density significantly increased in response to long-term (12 week) T+E2 treatment in WT but not in OPN-KO mice (C). Images of mouse prostate sections (4/tissue) were taken at 20x magnification. Col1a1 optical density (OD) was calculated via tissue segmentation to differentiate stroma from epithelium. One-way ANOVA was performed for each timepoint for (A). WT sham vs. WT T+E2: *: p≤0.05; ***: p≤0.001. WT T+E2 vs. OPN-KO sham: #: p≤0.05; ###: p≤0.001. WT. T+E2 vs. OPN-KO T+E2: $: p≤0.05; $$: p≤0.01. Significance for (B) and (C) was tested via pair-wise comparison with Mann-Whitney test. WT sham vs. WT T+E2 or OPN-KO sham vs. OPN-KO T+E2: *: p≤0.05; ***: p≤0.001. WT T+E2 vs. OPN-KO T+E2: #: p≤0.05.

To understand how prostatic pathology is involved in this process, we first determined the effect of loss of OPN function on prostatic growth and proliferation. Prostate weights at weeks two and six following pellet implantation significantly increased in both WT and OPN-KO mice, except that of the LP on week two, in which case prostate mass was significantly reduced in OPN-KO T+E2 mice (Fig. S5). Using Ki-67 staining; however, we determined that proliferation was increased in both WT and OPN-KO T+E2 mice, but there were significantly fewer Ki-67-stained cells in OPN-KO vs. WT VP two weeks after surgery (Fig. 5B). Intriguingly, steroid hormone imbalance did not change proliferation in the DP and reduced proliferation in OPN-KO mice (Figure S6). These results indicate that other processes, such as an increase in secretory products, may contribute to the overall increase in prostate weights in mouse prostate lobes, whereas the VP may be suited better to examine BPH as a proliferative disease.

Given our prior data showing strong association of extracellular matrix production with steroid hormone imbalance, and the growing recognition of benign prostatic hyperplasia as a fibrotic disease, we assessed the optical density of alpha-1 type I collagen subunit (COL1A1) as a marker of fibrosis using IHC. We screened the VP and DP for a significant change at two, six and twelve weeks after surgery (data not shown), and identified an increase only at week twelve in the VP. As it was expected based on the role of OPN in fibrotic diseases and our prior data on its pro-fibrotic role in bacterial prostatitis (16), we found that the loss of OPN significantly reduced COL1A1 protein accumulation in the VP (Fig. 5C).

## Discussion

Prostatic steroid hormone imbalance, described as the age-associated decline in testosterone and persistence of estradiol concentrations, has long been implicated in the pathogenesis of benign prostate disease (6–8). This is supported by rodent models of estradiol and testosterone hormone supplementation, that replicate human BPH most accurately to this date (9). Similar to human pathology, rodent models of steroid hormone imbalance recapitulate prostatic enlargement, proliferation and urinary dysfunction regulated via ERα estradiol receptor (17), however, mice do not develop nodular BPH as seen in humans. Inflammation and fibrosis are also accepted as components of benign prostate disease but the sequence of pathological events is uncertain (4, 18). Our study aimed to identify consequences of steroid hormone imbalance as they relate to early remodeling of the prostate immune environment and provoking long-term progression.

Our gene expression analysis identified the presence of major tissue remodeling in the VP with simultaneous extracellular matrix and degrading enzyme production indicating a dynamically changing tissue environment. Over time, with the progression of the model, matrix production likely becomes dominant over degradation fueling fibrosis as indicated by the elevated collagen I density after three months of steroid hormone imbalance. We also identified non-classical pro-inflammatory genes that were elevated by steroid hormone imbalance, namely *Saa1/2* and *Spp1*, whereas the expression level of most of the cytokines and chemokines on the Nanostring panel were below the level of detection.

We showed that *Spp1*+ macrophages infiltrated the glandular lumen because of steroid hormone imbalance. Furthermore, we demonstrated luminal macrophages to accumulate lipid droplets and acquire a foam cell phenotype in the prostates of mice with steroid hormone imbalance and in human BPH specimens. Our study is the first to report foam cell formation in benign prostate, which may be a particularly important step in the initiation of pathological changes. Foam cells have been implicated to drive inflammation and disease progression in atherosclerosis (19) and lung fibrosis (20). The infiltration of macrophages and their transition to foam cells drives the formation of the necrotic core of the plaque which eventually leads to the development of atherosclerotic lesions (19). The formation of foam cells in the prostate may occur due to augmentation of *de novo* lipid synthesis during steroid hormone dysregulation which is supported by our Nanostring analysis that identified an increase in *Fasn* expression, although, we did not identify the cellular source of lipid production in our study. Foam cells can form by the uptake of extracellular oxidized low-density lipoproteins (LDL) whereas hypoxia may also induce intrinsic lipid synthesis in macrophages (21). We also identified an overall decrease in genes encoding building blocks of the oxidative phosphorylation electron transport chain as well as disturbance in oxidative stress pathways. Mitochondrial dysfunction, defective oxidative phosphorylation and the accumulation of reactive oxygen species have been previously indicated to push metabolic pathways towards lipid synthesis (22, 23). The Nanostring panel, however, has its limitations on the availability of metabolic genes. Our findings clearly indicate that some form of lipid dysregulation occurs with steroid hormone imbalance, but future studies will be required to establish the cellular drivers and the conditions that led to the formation of foam cells.

Although foam cells have not been acknowledged in the context of BPH prior to our investigation, a few recent studies observed macrophages that contribute to lipid regulation in prostate cancer. One such study investigated tumor-associated macrophages (TAMs) and reported the presence of “lipid-loaded” macrophages that correlated with prostate cancer progression and stimulated prostate cancer cell migration via *Ccl6* secretion (24). Moreover, a recent study reported cholesterol-rich macrophages whose depletion reduced androgen signaling in prostate cancer (25). Foam cells in benign prostate, however, appear primarily in the luminal space from where their secreted messengers might reach a greater population of prostate cells. An indication for this wide range effect is the uptake of secreted osteopontin by prostate epithelial cells observed in our immunohistochemistry images. Possibly the closest example for the communication of luminal macrophages and neighboring cells is the pro-fibrotic action of lipid-laden macrophages in the lung, via the secretion of TGF-β (26). This indicates, that future investigations that assess the interactions of luminal macrophages with prostate epithelial cells are necessitated to advance our understanding on their role in BPH pathogenesis.

An intriguing feature of luminal macrophages is their ability to express and secrete high levels of osteopontin. On one hand, secreted osteopontin is expressed in various immune cells including macrophages, dendritic cells, neutrophils, eosinophils, NK cells, NKT cells, and T and B lymphocytes (27) whereas an alternatively translated intracellular form is expressed in macrophages and dendritic cells to drive pro-inflammatory gene expression and migration (28). Based on this, endogenous OPN may have contributed to the overall decrease in macrophage numbers in general as it was detected in all mouse prostate lobes. However, in the ventral prostate, which was the only lobe producing luminal macrophages, substantially more osteopontin was released shown via ISH, FISH and IHC. VP is also the lobe that have higher proliferation and fibrotic rate, immune cell and macrophage infiltration as well as, due to its anatomical proximity to the mouse prostatic urethra, it is most likely to affect urinary function. All of these aspects were improved in the absence of OPN which implies that luminal macrophage-derived OPN contributes to non-malignant pathologies in the prostate. It also highlights that the ventral prostate should be primarily assessed in investigations targeting macrophages in LUTD.

The overall increase in immune cells and tissue macrophages in response to steroid hormone imbalance, are also essential findings of our study. Tissue macrophages were most markedly elevated in the VP with a primarily M1 pro-inflammatory phenotype. This is particularly important, since previous studies provided conflicting evidence on macrophage densities in this model, which might have been due to the selection of later timepoints or other lobes as the target of investigation (13). Our results indicate an actively remodeling immune environment driven by macrophages in the VP. In a sharp contrast, mast cells are found in much higher numbers in the DP vs. the VP, with a significantly increased rate in steroid hormone imbalance and reduction by the loss of OPN. This may be partially explained by the ability of immobilized matrix OPN (but not soluble OPN), to retain mast cells (29). Increased degranulation in response to steroid hormone implants, was not observed, however, there was a significant decrease in OPN-KO T+E2 vs. WT T+E2 which is most likely attributable to the overall decrease in mast cell numbers, not reduced activation. This implies that changes in the steroid milieu promote mast cell retention, maybe through increased ECM deposition. However, they do not lead to increased mast cell activation, at least, early in pathogenesis. Mast cell numbers have been shown to increase in BPH and human pathogenic *E. coli* increases both the numbers and activation of mast cells in mice, in the dorsolateral lobe at the highest (30, 31). Although we saw no change in activation, it is plausible that increased mast cell numbers due to steroid hormone imbalance may predispose the prostate to evoke a higher mast cell response during inflammation. Although, we have insufficient evidence, these aspects should be investigated in the DP amongst other lobes.

Lobe-related diversity in responses to steroid hormone imbalance are plausible and the aspect of translational applicability of mouse prostate lobes has been the target of numerous investigations (32–34). Although a previous study using laser capture microdissection and microarray identified that the dorsal lateral lobe combination have closest similarities to the human peripheral prostate zone (32), a subsequent study using scRNA-seq determined that acinar cells are most similar in the lateral and ventral lobes (33). Also, stromal cell composition is altered across different lobes which feature has been implicated to drive the increased resistance to castration in the ventral lobe (34). Nevertheless, our study highlights that immunological responses are altered across the different mouse prostate lobes but also identifies the ventral lobe to gain pathological responses (inflammation, proliferation and collagen accumulation) most similar to human BPH during steroid hormone imbalance.

In summary, our study identified that steroid hormone imbalance increases macrophage numbers, drives their translocation to the lumen and their differentiation to lipid-laden foam cells. The pathological role of luminal macrophages is demonstrated by the action of secreted OPN on proliferation, fibrosis and urinary function. We also identified that retainment of mast cells in the DP is promoted by steroid hormone imbalance, which may sensitize immunological responses in the prostate.

## Materials and Methods

All materials and methods are available in the SI Appendix. Expression data is accessible through GEO Series accession number GSE220444.

## Acknowledgments

We thank Emily Ricke, Teresa Liu, Alexis Adrian, Hannah Miles, Han Zhang, Christian Ortiz Hernandez, Ana Lucila (Lulu) Bautista-Ruiz, Samuel Hubbard for experimental assistance and advice. The authors thank the University of Wisconsin Translational Research Initiatives in Pathology laboratory (TRIP) and the Carbone Cancer Center BioBank supported by the UW Department of Pathology and Laboratory Medicine, UWCCC (P30 CA014520) and the Office of The Director-NIH (S10 OD023526) for use of its facilities (slide scanning and NCounter assay) and services and for providing frozen human tissues. This research was funded by grants from the National Institutes of Health including K12 DK100022, K01 DK127150 and start-up fund from the Eastern Virginia Medical School(to P.P.), U54 DK104310 (to W.A.R. and C.M.V.), R01DK127081 and R01 DK131175 (to W.A.R).

## Notes

**Competing Interest Statement:** The authors declare no conflict of interest.

### Competing Interest Statement

The authors have declared no competing interest.

