## Supplemental materials for "Steroid hormone imbalance drives macrophage infiltration and *Spp1*/osteopontin^+^ foam cell differentiation in the prostate"

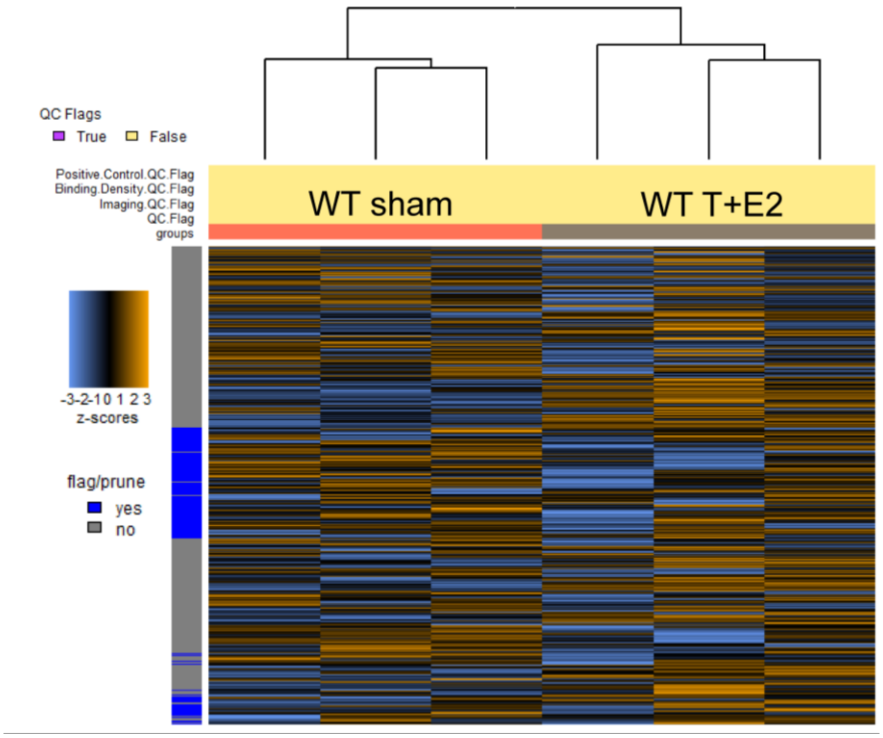


Figure S1: Heatmap shows a clear separation of samples into WT sham and WT T+E2 clusters based on normalized gene-expression. Gene-expression was analyzed using the nCounter mouse fibrosis panel from the ventral prostates of steroid hormone-implanted (estradiol and testosterone) or sham animals, two weeks after surgery. Heatmap was generated using the nCounter Advanced Analysis Software Genes flagged for low expression (blue in side column) were excluded from further analysis.

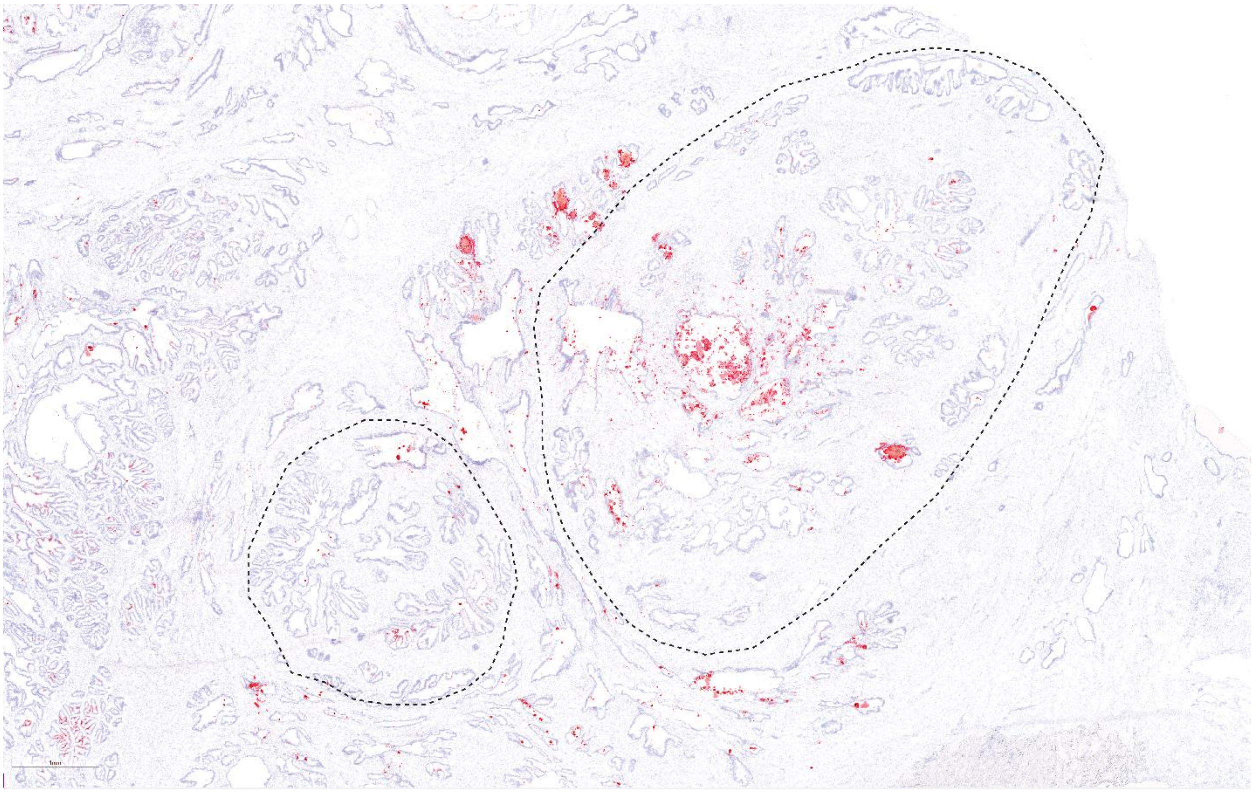

Figure S2: Lipid accumulation shows regionality in BPH tissues. Lipid content was stained with Oil Red O staining on a frozen BPH section. Slide was scanned at 40x magnification on a Leica Aperio AT2 slide scanner. Scale corresponds to 1mm.

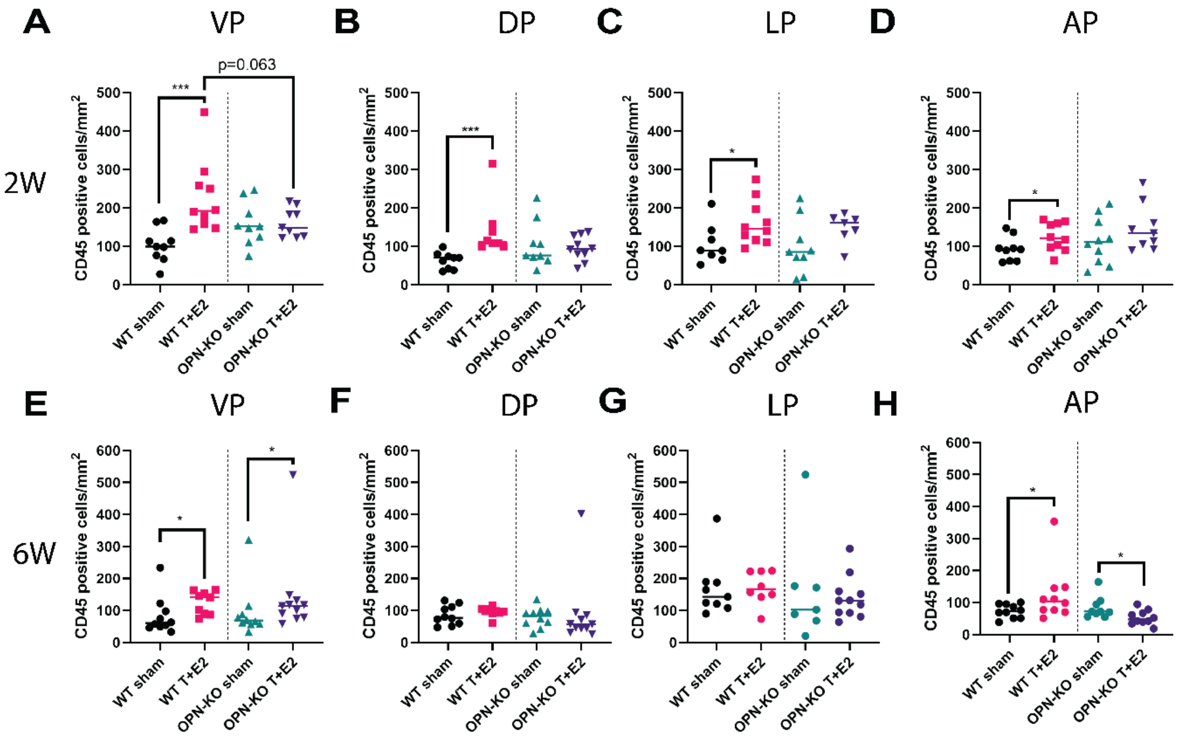


Figure S3: Immune cell numbers increase in all mouse prostate lobes in response to steroid hormone imbalance. CD45$^{+}$ pan-immune cell marker and immunohistochemistry were used to quantify immune cells in the ventral (VP, A and E), dorsal (DP, B and F), lateral (LP, C and G) and anterior (D and H) mouse prostate lobes two (2W, A, B, C and D) or six (6W, E, F, G and H) weeks after steroid hormone pellet implantation. Significance was tested by Mann-Whitney test. WT sham vs. WT T+E2 or OPN-KO sham vs. OPN-KO T+E2: * $p \leq 0.05$; *** $p \leq 0.001$.

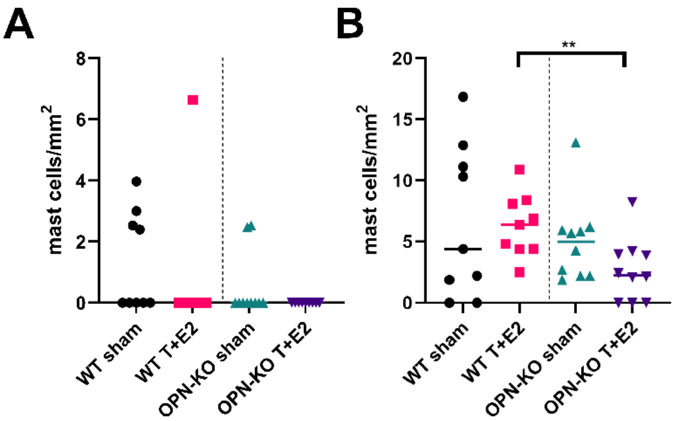


Figure S4: Activation of mast cells is unaltered by steroid hormone imbalance. Activated mast cells were counted in the ventral (A) and the dorsal (B) prostate based on the presence of released granules. Significance was tested by Mann-Whitney test. Slides were stained with a standard Toluidine blue staining method.

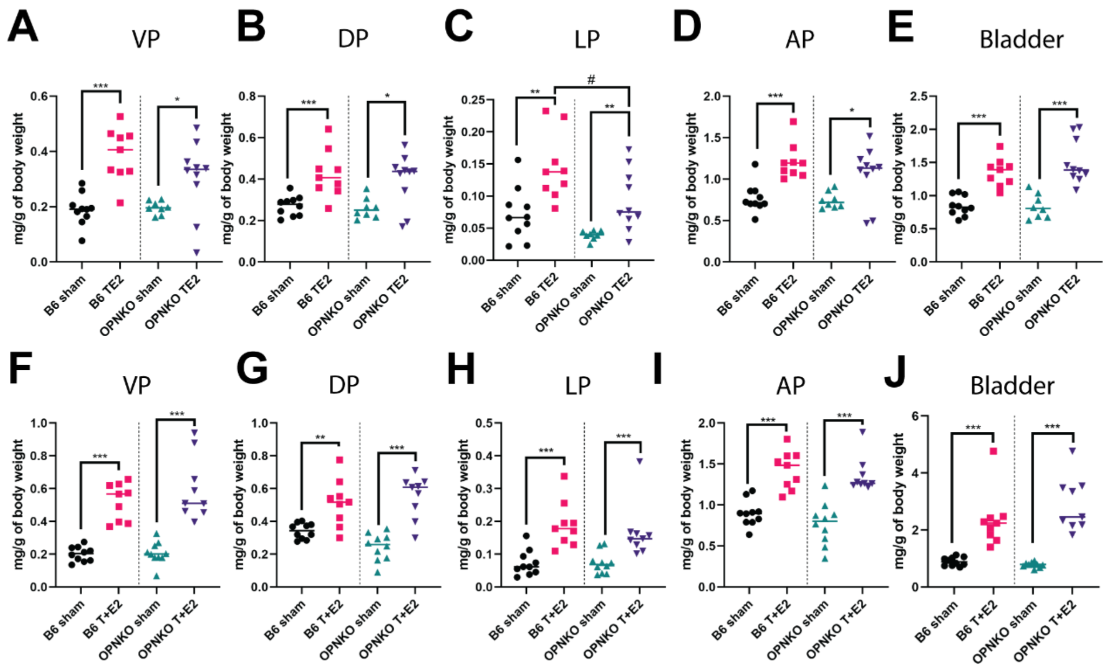


Figure S5: Overall, the loss of OPN did not substantially reduce steroid hormone-induced prostatic enlargement or bladder hypertrophy. The ventral (VP, A and F), dorsal (DP, B and G), lateral (LP, C and H) and anterior (D and I) mouse prostate lobes and the bladder (E and J) were weighed two (A, B, C, D and E) or six (F, G, H, I and J) weeks after steroid hormone pellet implantation. Only the LP mass was significantly reduced in OPN-KO mice at week two. Significance was tested by Mann-Whitney test. WT sham vs. WT T+E2 or OPN-KO sham vs. OPN-KO T+E2: *: $p \leq 0.05$; **: $p \leq 0.01$; ***: $p \leq 0.001$. WT T+E2 vs. OPN-KO T+E2: ##: $p \leq 0.01$.

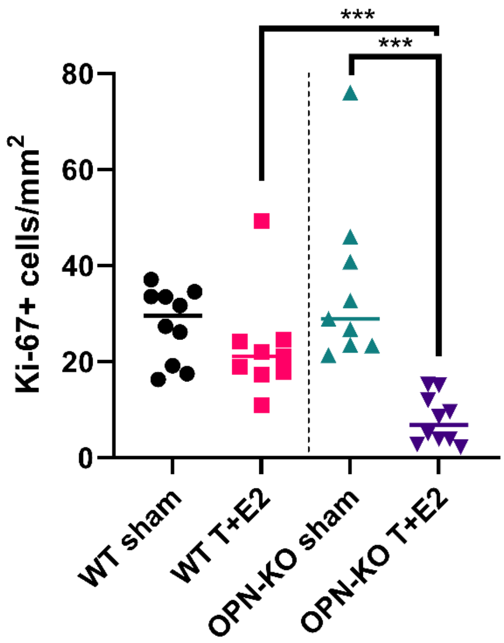


Figure S6: Steroid hormone imbalance did not affect proliferation in the dorsal prostate in WT animals. In contrast, it reduced proliferation in OPN-KO mice. Proliferation was measured by counting Ki-67$^+$ cells. Significance was tested by Mann-Whitney test. ***: $p \leq 0.001$

Supplementary methods

Ethics Statement

All experiments were conducted under approved protocols from the University of Wisconsin (ID: M005570-R01-A04) and the Eastern Virginia Medical School Animal Care and Use Committee (ID:22-011) and in accordance with the National Institutes of Health Guide for the Care and Use of Laboratory Animals.

Human tissues

Frozen tissues from patients with BPH were obtained via the University of Wisconsin Carbone Cancer Center BioBank and were selected by a pathologist. Five specimens were selected for immunohistochemistry and lipid staining and we selected examples based on visible nodule formation. No patient data, progression stage or symptom score was associated with the specimens.

Mice

Breeding pairs of Spp1tm1Blh/J and male C57BL/6J mice for wild type (WT) controls were obtained from The Jackson Laboratory. The Spp1tm1Blh/J strain was inbred at the UW Biomedical Research Model Services for no more than 3 generations. This strain was backcrossed to C57BL/6J for more than 10 generations by the donating lab, and the live colony was established in C57BL/6J mice using cryopreserved sperm at The Jackson Laboratory. Animals were maintained on a strict 12:12 h light-dark cycle in a temperature- and humidity-controlled facility with water and food provided *ad libitum*.

Void Spot Assay

Void spot assays (VSAs) were conducted as described previously (1). Briefly, mice were placed individually in cages lined with filter paper for 4 h with food but without water. Filter papers were imaged with an Autochemi AC1 Darkroom ultraviolet imaging cabinet (UVP). Images were imported into ImageJ, and void spots were quantified with Void Whizzard (2). Mice were acclimatized to the VSA conditions by performing the assay once without analyzing the data and then obtaining the baseline VSA parameters from the subsequent experiment. VSAs were repeated weekly for five weeks.

Hematoxylin-Eosin (H&E) and Toluidine blue staining and Immunohistochemistry (IHC)

H&E staining was performed by a standard method using Shandon Instant Hematoxylin and Eosin-Phloxine (Thermo Fisher Scientific) multichrome stain. Toluidine blue staining was also generated using a standard method with Toluidine blue O dissolved in 70% ethanol. For IHC, sections were de-paraffinized and hydrated. Antigen retrieval was acquired in a decloaking chamber (Biocare Medical) using citrate buffer pH 6.0. Endogenous peroxidases and non-specific binding sites were blocked with Bloxall (Vector Laboratories) and horse serum (2.5% or 10%) or Rodent Block, respectively. Primary antibodies anti-Col1a1 (NBP30054, Novus Biologicals,1:300 dilution), anti-CD45 (ab10558, Abcam, 1:2000 dilution), anti-Vimentin (bs-0756R, Bioss, 1:300 dilution), anti-Ki-67 (VP-K452, Vector Laboratories, 1:1000 dilution) and anti-OPN (AF808, R&D Systems, 1:100 dilution) were added overnight (Human CD68). An HRP-conjugated horse anti-rabbit or anti-mouse IgG Polymer (Vector Laboratories) or rabbit anti-goat IgG antibody (Bethyl Laboratories, 1:250 dilution) was used as secondary for 30 min and the signal was developed using the SignalStain DAB Substrate Kit (Cell Signaling).

Oil Red O staining and IHC on frozen sections

Frozen prostate sections were briefly air-dried and were fixed in 4% formaldehyde in PBS for 10 minutes. Sections were briefly washed with tap water and stained with freshly prepared Oil Red O staining (prepared in isopropanol and water). Sections were rinsed with 60% isopropanol and stained with Mayer`s hematoxylin. Slides were mounted with aqueous mounting medium. IHC with frozen sections started with a 15-minute air-dry, a 10-minute fixation in 10% neutral buffered formaldehyde and included no antigen retrieval. Slides followed the regular IHC protocol detailed above after the antigen retrieval step. An anti-CD68 antibody (ab213363, Abcam, 1:8000 dilution) was applied overnight to label macrophages.

### RNAscope and Fluorescent in situ hybridization

*In situ* hybridization was performed with the RNAscope method using probe sets specific to mouse Spp1 (435191, target region: 2–1079 bp,) and Mmp12 (406551, target region: 1949–2956) with the RNAscope® 2.5 HD Assay-Red or -Brown detection system (Advanced Cell Diagnostics). Reaction specificity was tested with a positive control probe and a negative control with only a probe diluent instead of the probe. Fluorescent in situ hybridization (FISH) was performed using the Hybridization Chain Reaction (HCR) technique from Molecular Instruments with probes targeting Spp1, Cd68, Cd3e, Cd19 and Saa1/2 according to the manufacturer`s instructions.

### Imaging and Analysis

Tissues were imaged with a Mantra 2 Quantitative Pathology Workstation (Akoya Biosciences) with a 20x (for Col1a1) or a 40x objective (for all others). Four (for 20x) or six (for 40x) representative images were taken per tissue. Area not containing tissue was removed, or only the stroma was selected (for Col1a1 density) to acquire the total tissue area for normalization. Optical density (OD) or % positivity was quantified by inForm software (Akoya Biosciences). Number of positive cells for Ki-67 and all immune markers were counted by hand. Cell counts were normalized to tissue area.

All four experimental groups were represented on each tissue blocks/slides, and slides from the same tissue and timepoint were processed together.

Human prostate tissues were also imaged by a slide scanner at 40x magnification on a Leica Aperio AT2 slide scanner.

### RNA Isolation and Nanostring analysis

Ventral lobe RNA was stabilized in RNAlater (Thermo Fisher Scientific) and total RNA was isolated from ventral lobes using the Qiagen RNeasy Micro kit with on-column DNAse digestion (Germantown, MD, USA). RNA quantity and quality was measured with a NanoDrop. RNA expression was analyzed with the nCounter mouse fibrosis panel (NS_Mm_Fibrosis_V2.0). Analysis of differentially expressed genes, cell-type enrichment and gene set analysis was performed using the nCounter Advanced Analysis Software. Volcano plots were generated using VolcaNoseR (3).

### Statistical Analysis

All statistical calculations were performed in Graphpad Prism (Graphpad Software, San Diego, CA, USA). One-way ANOVA was used if the Brown–Forsythe and Bartlett's tests were significant followed by Tukey's post hoc analysis. Otherwise, the non-parametric Kruskal–Wallis test with Dunn's multiple comparison test for groups more than 2 was performed. To compare two groups, we used two-tailed t-test when the F-test was significant. Otherwise, the statistical difference was determined by Mann–Whitney test.
